# Structures of FHOD1-Nesprin1/2 complexes reveal alternate binding modes for the FH3 domain of formins

**DOI:** 10.1101/2020.06.19.161299

**Authors:** Sing Mei Lim, Victor E. Cruz, Susumu Antoku, Gregg G. Gundersen, Thomas U. Schwartz

## Abstract

The nuclear position in eukaryotic cells is controlled by a nucleo-cytoskeletal network, with important roles in cell differentiation, division and movement. Forces are transmitted through conserved linker of nucleoskeleton and cytoskeleton (LINC) complexes that traverse the nuclear envelope and engage on either side of the membrane with diverse binding partners. Nesprin-2 giant (Nes2G), a LINC element in the outer nuclear membrane, connects to the actin network directly as well as through FHOD1, a formin whose major activity is bundling actin. Much of the molecular details of this process remain poorly understood. Here, we report the crystal structure of Nes2G bound to FHOD1. We show that the G-binding domain of FHOD1 is rather a spectrin repeat binding enhancer for the neighboring FH3 domain, possibly establishing a common binding mode among this subclass of formins. The FHOD1-Nes2G complex structure suggests that spectrin repeat binding by FHOD1 is likely not regulated by the DAD helix of FHOD1. Finally, we establish that Nes1G also has one FHOD1 binding spectrin repeat, indicating that these abundant, giant Nesprins have overlapping functions in actin-bundle recruitment for nuclear movement.

## INTRODUCTION

Eukaryotic cells in multicellular organisms display an enormous range of specialization. The development of different tissues coincides with cellular reorganization, particularly of the elaborate organellar structure. The position of the nucleus as the largest organelle is often found to be a distinct marker for certain cell types, for example in muscle cells or in neurons^1^. During nuclear migration, the nucleus interacts with the cytoskeletal network for active positioning. Pulling forces across the nuclear envelope are mediated by a two-protein complex known as Linker of Nucleoskeleton and Cytoskeleton (LINC)^2,3^. LINC complexes are universally conserved in eukaryotes and consists of Nesprins (Nuclear envelope spectrin repeat proteins) that transverse the outer nuclear membrane (ONM), and Sad1-UNC84 (SUN) proteins which pinch through the inner-nuclear membrane (INM) and connect to the nucleoskeleton. In the perinuclear space, three SUN domains interact with three C-terminal KASH peptides of Nesprins to form an intricate heterohexameric assembly^4^, the core of the LINC complex. Humans have five known SUN proteins and six KASH-containing ONM proteins. Some of these SUN- and KASH-proteins are ubiquitously expressed, while some are tissue specific. They generate a diverse network of nucleo-cytoplasmic linkages that are important for homeostasis and trigger multiple genetic diseases, if altered ^5–7^. The regulation and the interplay between the different LINC complexes is an important element in deciphering the entire network. In the migrating fibroblasts, the nuclear position depends on nesprins-2 giant (Nes2G), a large, actin-binding LINC component of transmembrane actin-associated (TAN) lines^8^. FHOD1 is a formin that itself binds and bundles actin, while also interacting with Nes2G^9^. This way, an interaction network between actin bundles, Nes2G, and FHOD1 is established.

The molecular details of this intricate connection are largely unknown, in no small part because the proteins involved are complicated, multi-domain entities. The 800 kDa Nes2G contains two N-terminal, actin-interacting calponin homology (CH) domains followed by 56 spectrin repeats (SRs) before the ONM-transversing transmembrane-helix and the C-terminal KASH peptide, that interacts with SUN1/2^10,11^. Formins are categorized into several classes dependent on their function and domain architecture^12–15^. FHOD1 is the founding member of one class, with an N-terminal, presumed G-protein binding domain (G2 or GBD2), followed by an FH3 or diaphanous inhibitory domain (DID) that is autoinhibited through a conserved C-terminal diaphanous-autoregulatory domain (DAD)^16^. Between FH3 and DAD, FHOD1 also contains a dimerization motif and the actin-bundling FH2 domain, central to all formins^12,17^. Another large class of formins are the diaphanous-related formins (DRFs). They contain a tested, structurally different GBD N-terminal to the FH3^18,19^.

We sought to advance our mechanistic understanding of the emerging, functionally important actin-Nes2G-FHOD1 network. In this study, we determined the crystal structure of Nes2G- SR11/12 in complex with FHOD1, revealing a novel formin-binding motif within spectrin repeats. Using a bioinformatic analysis we detect that Nes1G also carries this formin-binding motif in one out of its 76 SRs, which we confirm with structural and biochemical methods. SR11/12 binding by FHOD1 is outside of the autoregulation through DAD and independent of actin bundling. Further, we establish that the small domain preceding the FH3 domain of FHOD1, formerly GBD2 or G2, is in fact a modulator of the binding specificity of the neighboring FH3 domain, making the paired domain specific for SR, rather than GTPase interaction.

## RESULTS

### Crystal structure of Nesprin2G-SR11/12 in complex with FHOD1-N

We set out to characterize how Nes2G interacts with the N-terminal GBD-FH3 domain element of FHOD1 structurally. This is based on a previous study where specific Nes2G SRs and FHOD1 domain domains have been identified^9^. We recombinantly expressed spectrin repeats SR 11-12 of human Nes2G (residues 1425-1649) as well as FHOD1-N (residues 1-339) (Figure 1a)^20^. Both proteins form a stable 1:1 complex, with a measured K_D_ of 375 nM (Supplementary fig. 1). While we initially worked with a cysteine double-mutant, FHOD1-N (C31S C71S), based on the report that wildtype FHOD1-N have a tendency to form artificial, cysteine-mediated dimers in solution^20^. We did not observe the reported behavior (Supplementary fig. 2). Therefore, we continued our studies with wildtype FHOD1-N.

**Table 1.** Data collection and refinement statistics (molecular replacement)

|  | Human Nes2G-SR11/12<sub>1425-1649</sub> | Human Nes1G-SR17/18<sub>1976-2200</sub> |
| --- | --- | --- |
|  | FHOD1<sub>1-339</sub> | FHOD1<sub>1-339</sub> |
| **Data collection** |  |  |
| Space group | C 2 | C 2 |
| **Cell dimensions** |  |  |
| *a*, *b*, *c* (Å) | 237.8, 52.4, 117.5 | 245.1, 57.7, 140.8 |
| $\alpha$, $\beta$, $\gamma$ (°) | 90.0, 123.3, 90.0 | 90.0, 125.5, 90.0 |
| Resolution (Å) | 67.3 - 2.8 (2.9 - 2.8) | 99.0 - 7.0 (7.3 - 7.0) |
| $R_{\text{pim}}$ | 0.07 (1.3) | 0.16 (0.9) |
| *I* / $\sigma(I)$ | 21.6 (1.9) | 6.1 (0.8) |
| $CC_{1/2}$ | 0.54 | 0.50 |
| Completeness (%) | 98.9 (96.1) | 97.7(91.5) |
| Total reflections | 50981 | 5293 |
| Unique reflections | 8641 | 2635 |
| Redundancy | 5.9 | 2.0 |
| **Refinement** |  |  |
| Resolution (Å) | 67.3 - 2.8 | 99.7 - 7.0 |
| No. reflections | 45282 | 2634 |
| $R_{\text{work}}$ / $R_{\text{free}}$ | 0.22 / 0.27 | 0.37 / 0.42 |
| **No. atoms** | 8568 | 6062 |
| Protein | 8536 | 6062 |
| Ligand/ion | 10 |  |
| Water | 22 |  |
| **B-factors (Å<sup>2</sup>)** |  |  |
| Protein | 77.5 | 400.5 |
| Ligands/ion | 112 |  |
| Water | 80.4 |  |
| **r.m.s. deviations** |  |  |
| Bond lengths (Å) | 0.01 | 0.02 |
| Bond angles (°) | 1.5/ | 2.3 |
| **Ramachandran analysis** |  |  |
| Favored (%) | 96.3 | 96.4 |
| Allowed (%) | 3.3 | 3.5 |
| Outliers (%) | 0.4 | 0.1 |

\*Values in parentheses are for highest-resolution shell.

**Figure 1:**
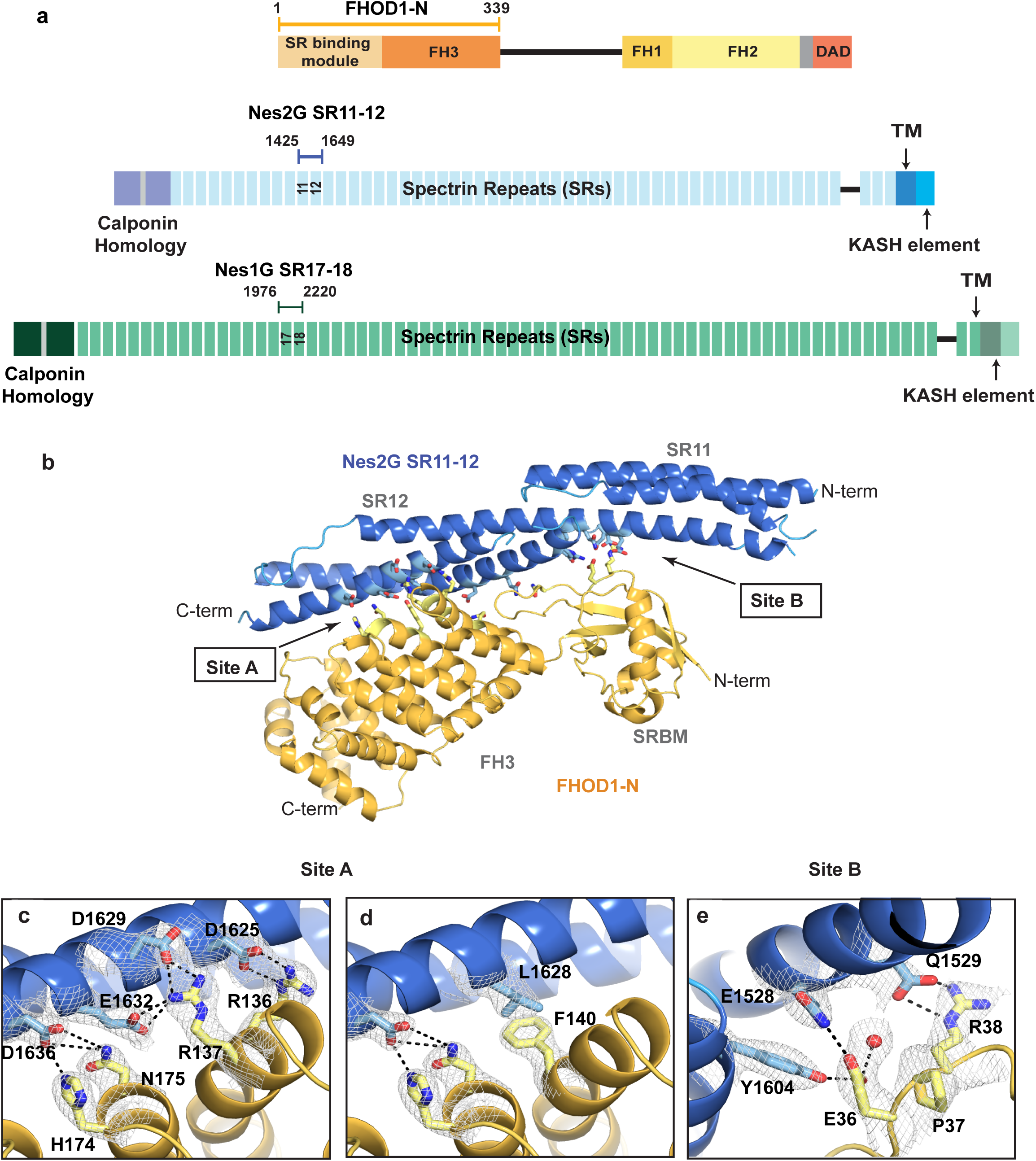
Structure of the FHOD1-N and Nes2G-SR11/12 complex. **a**, Schematic diagram to show the boundaries of FHOD1-N, Nes2G-SR11/12, and Nes1G- SR17/18 fragments in the context of the full-length protein. **b**, Overall crystal structure of Nes2G-SR11/12 bound to FHOD1-N with interface residues, as part of the main interaction sites A and B shown as sticks. Nes2G in blue, FHOD1 in orange. **c, d**, Close-up views of key interaction sites within site A. C and D show the same perspective, with different, otherwise overlapping residues, removed for clarity. H-bonds and charged interactions shown as black dashed lines. **c**, emphasizes charged interactions, **d**, a critical hydrophobic interaction. **e**, Close- up view of site B dominated by the EPR_36-38_ element of the SRBM of FHOD1. A water molecule in red.

**Figure 2:**
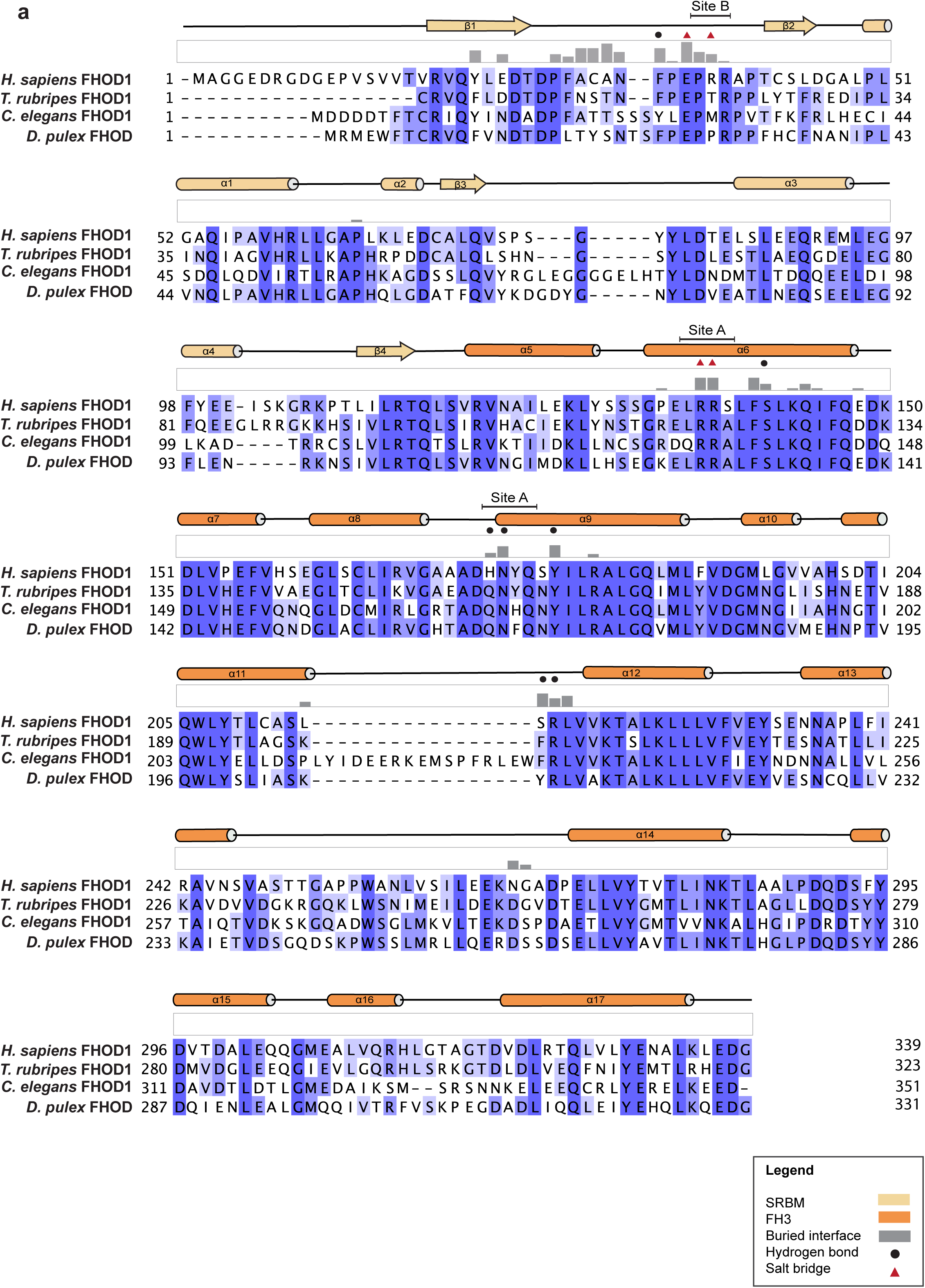
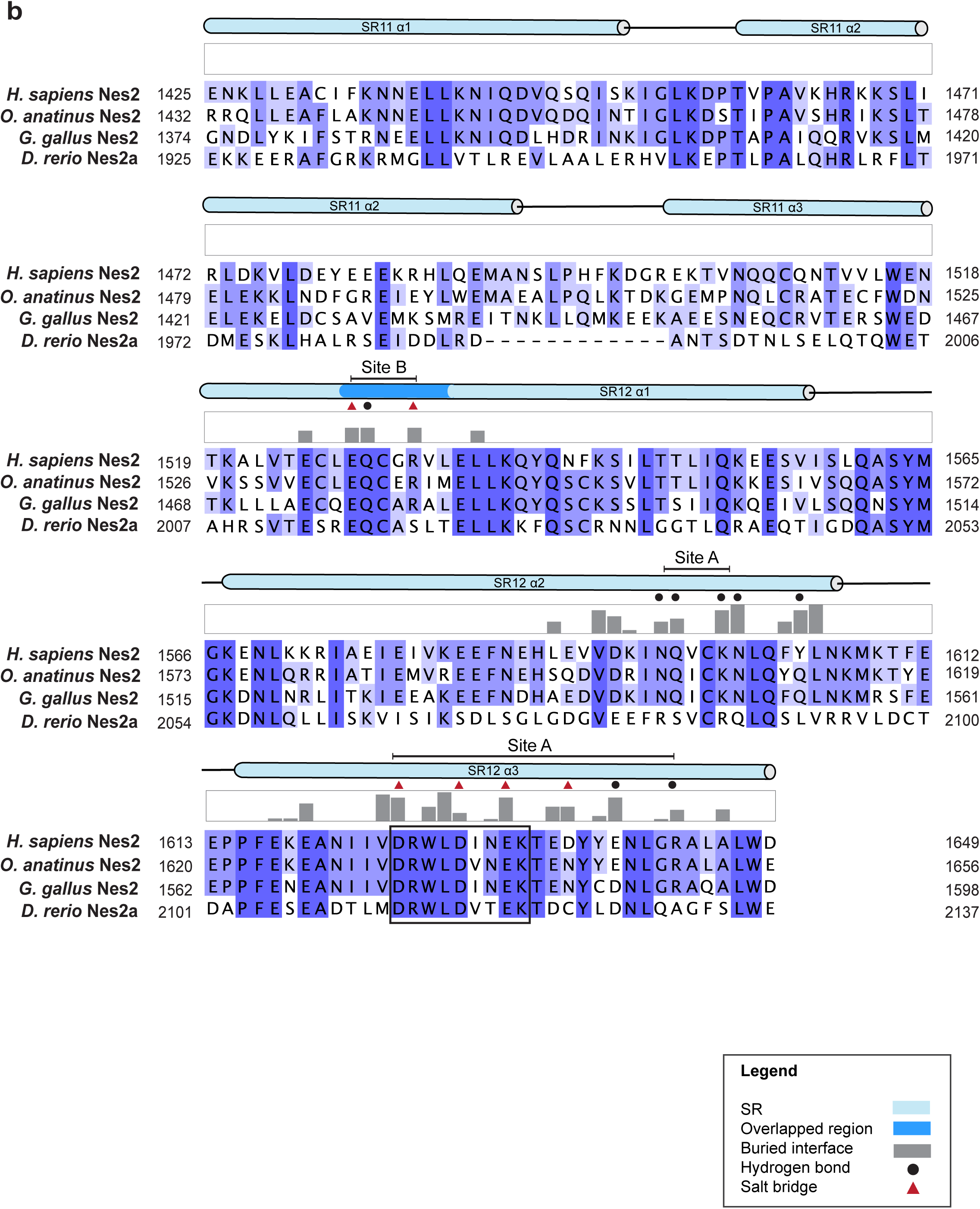
Sequence alignment and conservation analysis of FHOD1-N and Nes2G-SR11/12. **a**, Sequence alignment for FHOD1-N and **b**, Nes2G-SR11/12 using a blue-to-white color gradient to indicate high-to-low conservation. Secondary structure elements are indicated above the sequence alignment. Interface residues of the complex interaction sites are indicated by grey bars, representing the degree of burial upon binding. Other symbols denote salt bridges (red triangles) and hydrogen-bonds (black discs).

We obtained crystals of the Nes2G-SR11/12-FHOD1-N that diffracted to 2.8 Å resolution. The crystals belong to space group C2 and contain two heterodimeric complexes per asymmetric unit. One of the two complexes is better packed and therefore better resolved. Both complexes superpose well with a root mean square deviation (RMSD) of 0.82 Å. Nes2G-SR11/12 adopts the canonical structure of a tandem spectrin repeat (SR), i.e. two antiparallel coiled-coils connected by a continuous α-helix, generating two tethered three-helix bundles with a left-handed twist (Figure 1b). FHOD1-N superposes well with the apo-structure^20^ with an RMSD of 1.08 Å. The FH3 domain (residues 115-339) is an α-helical solenoid composed of five armadillo repeats. It is immediately preceded by a small domain (residues 14-114) that has a ubiquitin superfold. Formerly described as a GTPase-binding domain, we refer to this domain as an SR-binding module (SRBM), as it will become clear further on. The most significant difference between apo- and Nes2G-bound FHOD1-N is the ordering of a surface loop in the SRBM domain, residues 30-40, upon binding.

FHOD1-N engages with the tandem SR repeat by generating a continuous binding surface along the long axis of the helical bundle. In total, the binding interface buries 1191 Å^2^. The majority of the interaction is between the FH3 domain and SR12, with additional contributions from the SRBM and the long, SR11/SR12 connecting central helix. As it is typical for protein- protein interactions, we see a mix of van der Waals, polar and charged interactions throughout the interface. Two areas within the binding interface stand out, referred to as site A and site B (Figure 1 b - e). Site A is the core interface, centered around a remarkable charge network. D1625, D1629, and E1632 of Nes2G-SR12 form a web of salt bridges with R136 and R137 of FHOD1-N (Figure 1c). The hydrophobic residues L1628 of Nes2G-SR12 and F140 of FHOD1-N get deeply buried in the site A interface upon binding. A number of additional hydrogen-bonds surround and complete this site (Figure 2). All the key residues in site A are very well conserved across diverse species, supporting the notion that this is an important functional interface (Figure 2). Site B involves the loop connecting strands β1 and β2 of FHOD1-SRBM, particularly residues 30-38, interacting primarily with the very N-terminal α-helical element Nes2G-SR12 (residues 1525-1532). The FHOD1-SRBM-loop packs against the SR12 helical bundle and becomes well-ordered, in contrast to its more flexible position in the apo-form. A hydrogen-bonding / charge network is formed between E36 and R38 of FHOD1-SRBM and E1528 and Q1529 of Nes2G- SR12, with the best ordered water molecule of the structure as an integral component (Figure 1e). Several exposed, hydrophobic residues get buried upon binding (A32, F34 of FHOD1- SRBM and F1603 of Nes2G-SR12) (Figure 2).

### Mutations in Nes2G-FHOD1 interface residues abolish protein complex formation

In order to determine which residues are most critical for the interaction, and to generate meaningful point-mutations for biological assays, we analyzed the effect of mutating conserved interacting residues. We probed for complex formation by size exclusion chromatography (SEC) and bio-layer interferometry (BLI) (Figure 3). In site A, we determined that the charge network is essential for binding, since a Nes2G-SR11/12 D1625A/D1629A double mutant abolished binding in the SEC assay. Mutating the paired, charged residues on FHOD1-N, R136A and R137A, expectedly, has the same effect (Figure 3b - d). The hydrophobic interaction between Nes2G-SR11/12 L1628 and FHOD1-N F140 is also essential. A Nes2G-SR11/12 L1628R or Nes2G-SR11/12 L1628A/E1632A mutant abolishes binding as well (Figure 3a & c). We also analyzed site B, probing it with a FHOD1-N SRBM-loop triple mutant E36A/P37G/R38A. SEC and BLI both still show residual binding, suggesting that site B is perhaps somewhat less critical than site A (Figure 3b - d). In summary, we establish Nes2G-D1625A/D1629A and FHOD1- R136A/R137A as structurally defined, non-interacting probes for *in vitro* and *in vitro* studies.

**Figure 3:**
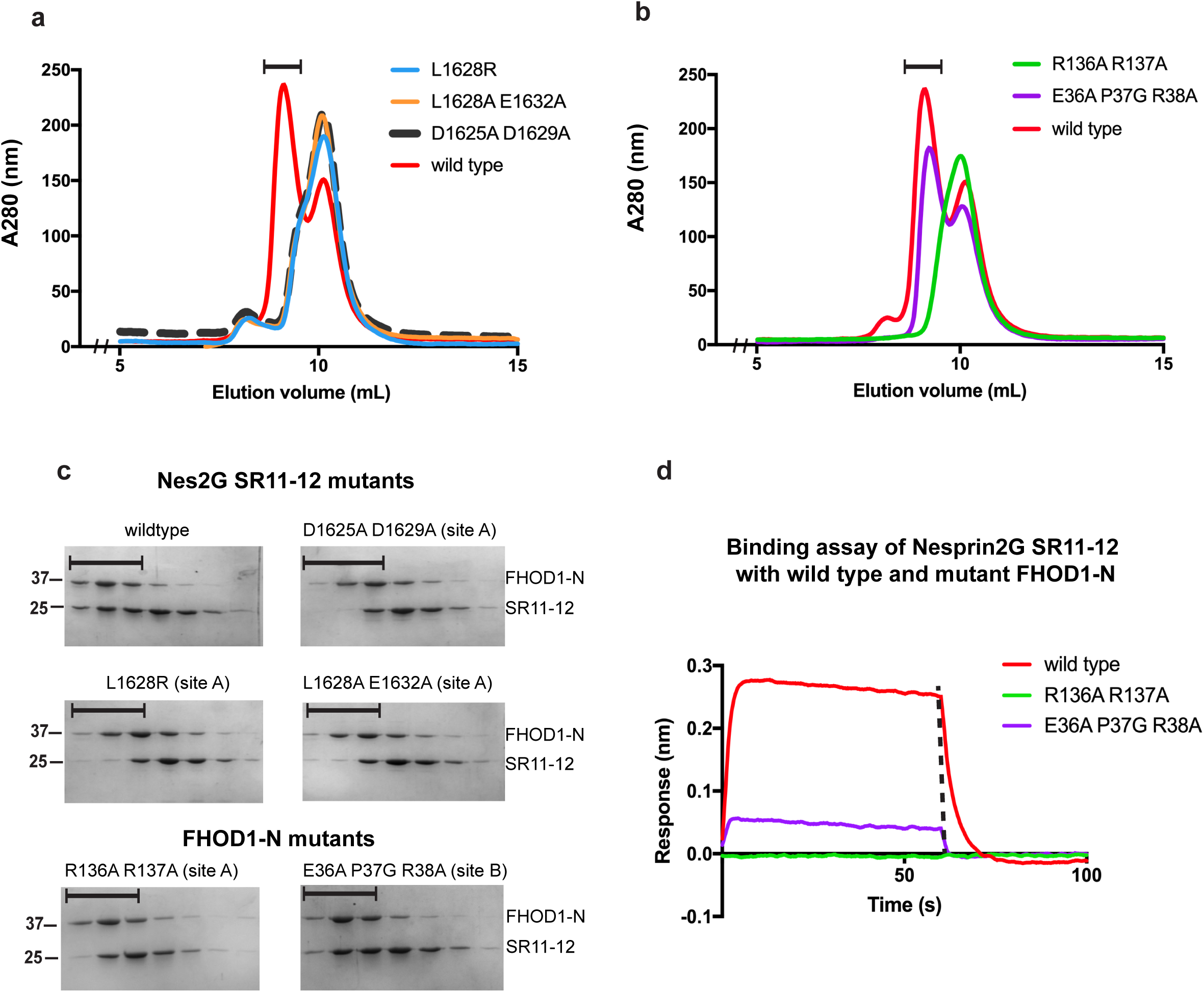
Analysis of residues involved in FHOD1-N-Nes2G-SR11/12 complex formation. **a**, Size exclusion chromatography (SEC) profile of purified FHOD1-N incubated with wildtype Nes2G-SR11/12, or mutant variants. Only the wildtype stably binds. **b**, Complementary experiment with Nes2G-SR11/12, and wildtype or mutant FHOD1-N fragments. Both experiments were carried out on a Superdex S75 10/300 column. **c**, SDS-PAGE analysis of the SEC elution fractions (black bars) in A and B. **d**, Binding activity of the two proteins with Biolayer interferometry (BLI) where Nes2G-SR11/12 is immobilized as ligand tested against the indicated analytes.

### FHOD1-mutations in Nes2G interface block nuclear migration

It is known that FHOD1 and Nes2G functionally interact in the context of nuclear migration^21,22^. We tested the FHOD1-R136A/R137A mutant in a nuclear movement assay. In the assay, wounded monolayers of serum-starved fibroblast cells are stimulated by lysophosphatidic acid (LPA), resulting in rearward movement of the nucleus away from the wounded cell edge. We depleted endogenous FHOD1 in NIH3T3 fibroblast using a shRNA approach, which resulted in nuclear movement defects that can be rescued by the expression of wildtype FHOD1 (Figure 4a - c). In contrast, we see hardly any rescue using the FHOD1-R136A/R137A. The result is a phenocopy of FHOD1-I705A mutant which inactivates actin barbed-end binding^23^. This experiment demonstrated that the direct FHOD1-Nes2G contact is critical for nuclear migration.

**Figure 4:**
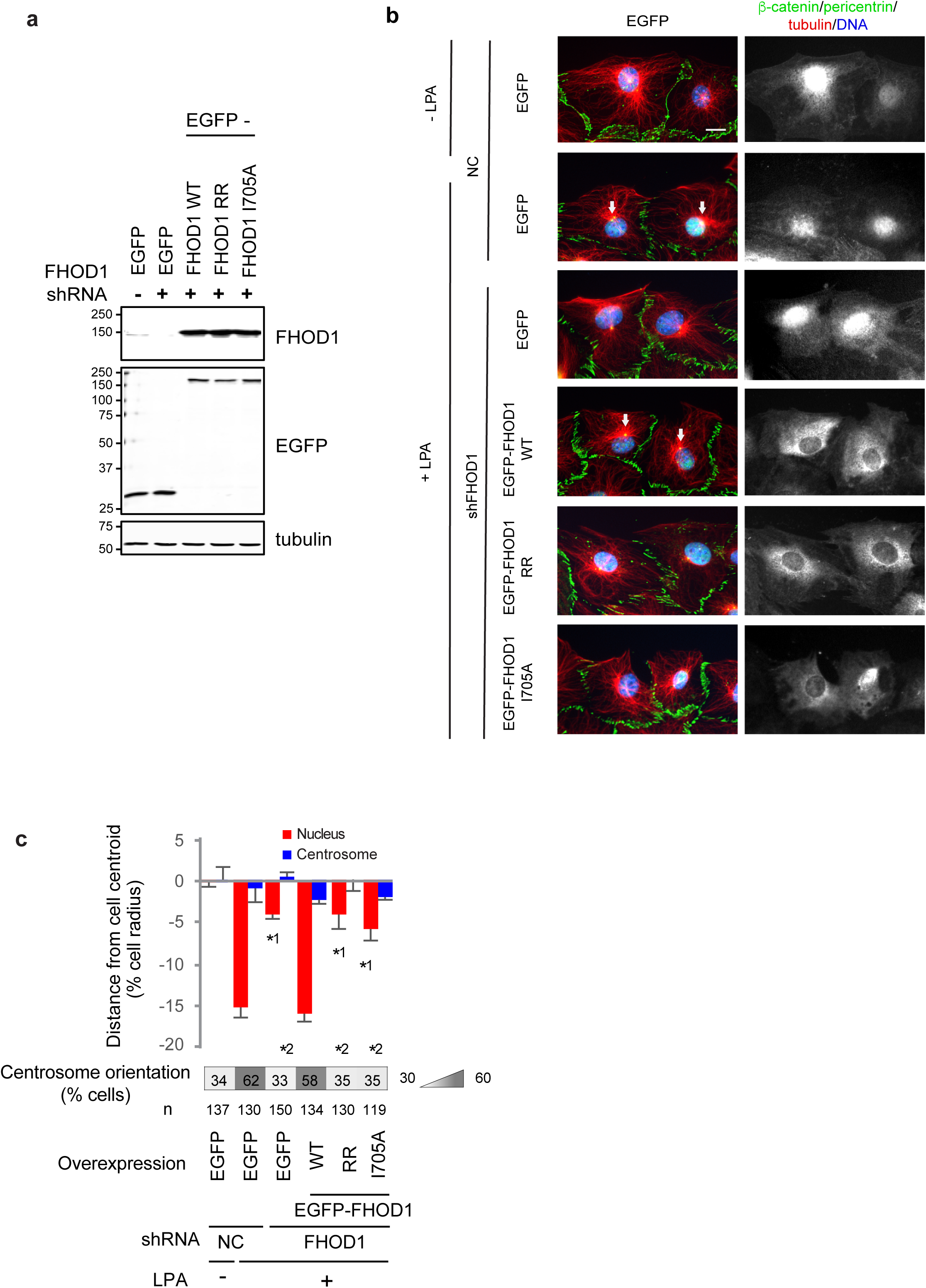
FHOD1 R136A R137A double mutant (RR) fails to rescue centrosome orientation and nuclear movement. **a**, Western blots of lysates from NIH3T3 fibroblasts with or without FHOD1 shRNA (shFHOD1) expression and expressing either EGFP or EGFP-tagged FHOD1 proteins. Antibodies used to probe the blots are indicated on the right. FHOD1 RR is FHOD1 with R136A R137A point mutations. **b**, Images of LPA-stimulated wound-edge NIH3T3 fibroblasts expressing EGFP or FHOD1 proteins after knockdown of FHOD1 (shFHOD1) and stained for the indicated proteins and DAPI. NC is a negative control shRNA. White arrows, oriented centrosomes. Bar, 20 μm. **c**, Quantification of centrosome and nuclear positions and centrosome orientation for the cells treated as in A. Values are means ± SEM; n, cells examined. Centrosome orientation (mean % of cells), is shown in the heat map below the histograms. *^1^ and *^2^ indicate p < 0.01 compared to the LPA-stimulated control to the rest of the samples for each category

### The FHOD1 binding motif is also found in Nesprin1-Giant

Having established how FHOD1-N interacts with Nes2G-SR11/12 we asked whether the motif we identified might be found in other spectrin repeat proteins as well. Based on our mutational study, we identified DxWLD[IVLA]xE to be a signature motif within a spectrin repeat that would suggest FHOD1-binding. By using BLAST pattern search, we found this motif in one other spectrin repeat containing actin binding protein, namely Nesprin1-Giant (Nes1G). Nes1G is also part of LINC complexes, with a C-terminal KASH-peptide that interacts with Sun1/2. Similar to Nes2G, the N terminus of Nes1G has two actin-binding CH domains followed by many (74 vs 56) SRs (Duong et al., 2014). We found the motif, 100% conserved, in Nes1G-SR18 (Figure 5a). To test the functionality of Nes1G-SR18 as an FHOD-1 interactor, we expressed and purified the tandem SR repeat Nes1G-SR17/18 (residues 1976-2200) recombinantly in *E. coli* and analyzed FHOD1-N binding by SEC and BLI. FHOD1-N and Nes1G-SR17/18 coelute from a SEC column as a complex with 1:1 stoichiometry (Figure 5b). We measured a binding affinity between Nes1G-SR17/18 and FHOD1-N of 85 nM, comparable to that of Nes2G-SR11/12 under the same buffer condition (Figure 5c). Further pointing to the same binding interaction, when the aspartate residues were mutated in the DxWLD[IVLA]xE motif or the corresponding R136A/R137A mutations introduced into FHOD1-N, Nes1G-SR17/18 no longer binds to FHOD1-N (Figure 5d, e).

**Figure 5:**
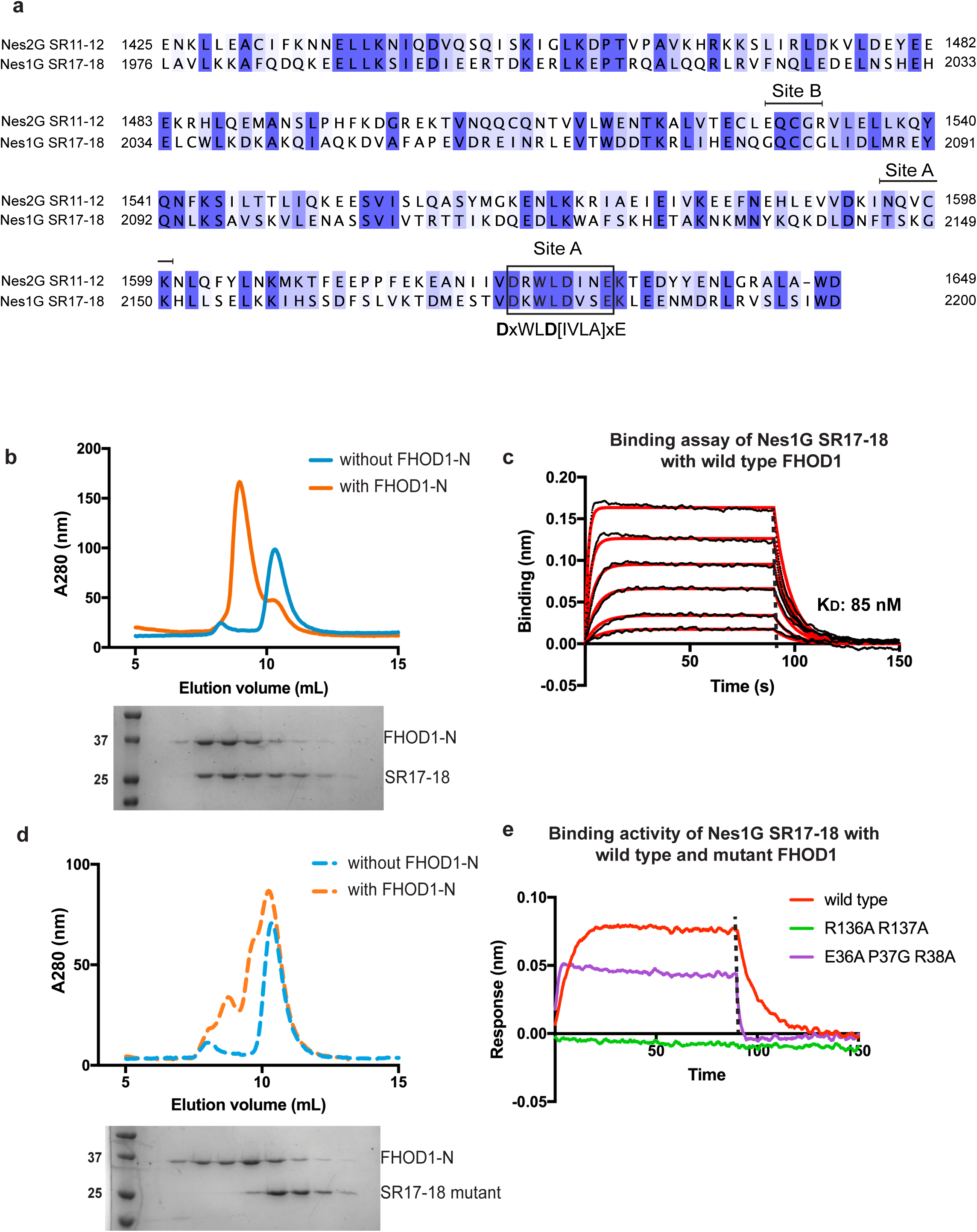
Nes1G-SR17/18 is another FHOD1-binding spectrin repeat. **a**, Sequence alignment of the two tandem SRs that have a matching sequence pattern indicative of FHOD1 binding. The search sequence is shown below the alignment. Conserved interface residues involved in site A and site B are labelled and the critical motif highlighted in box **b**, SEC profile of Nes1G-SR17/18 incubated with or without FHOD1-N, with SDS PAGE analysis of the eluted fractions. **c**, BLI binding assay of Nes1G-SR17/18 and FHOD1-N. Individual curves represent a 2-fold dilution series from a maximal concentration of 400 nM **d**, Setup as in **b** but using mutant Nes1G-SR17/18 D2176A D2180A instead of wildtype. Both SEC experiments were performed on a Superdex S75 10/300 column. **e**, Binding assay of wildtype and mutant FHOD1-N with Nes1G-SR17/18 measured by BLI.

### Structure of Nes1G-SR17/18 with FHOD1-N

We obtained crystals of Nes1G-SR17/18 in complex with FHOD1-N that diffracted to 7 Å resolution. The structure was solved by molecular replacement using the Nes2G-SR11/12- FHOD1-N complex as the search model. The crystals also belong to the C2 space group, with two complexes per asymmetric unit. Due to the modest resolution, we only used rigid body refinement to position the four domain elements, Nes1G-SR17, Nes1G-SR18, FHOD1-SRBM, and FHOD1-FH3. Nes1G-SR17 is essentially invisible in the density, suggesting disorder (Figure 6a, Supplementary fig. 3). The other three domain element superpose very well with the Nes2G-SR12-FHOD1-N complex. In order to verify whether SR17 is dispensable for FHOD1-N interaction, we analyzed a Nes1G-SR18-only construct (residues 2076-2200) for FHOD1-N binding. We found stable FHOD1-N interaction with Nes1G-SR18 as analyzed by SEC and BLI (Figure 6 b,c).

**Figure 6:**
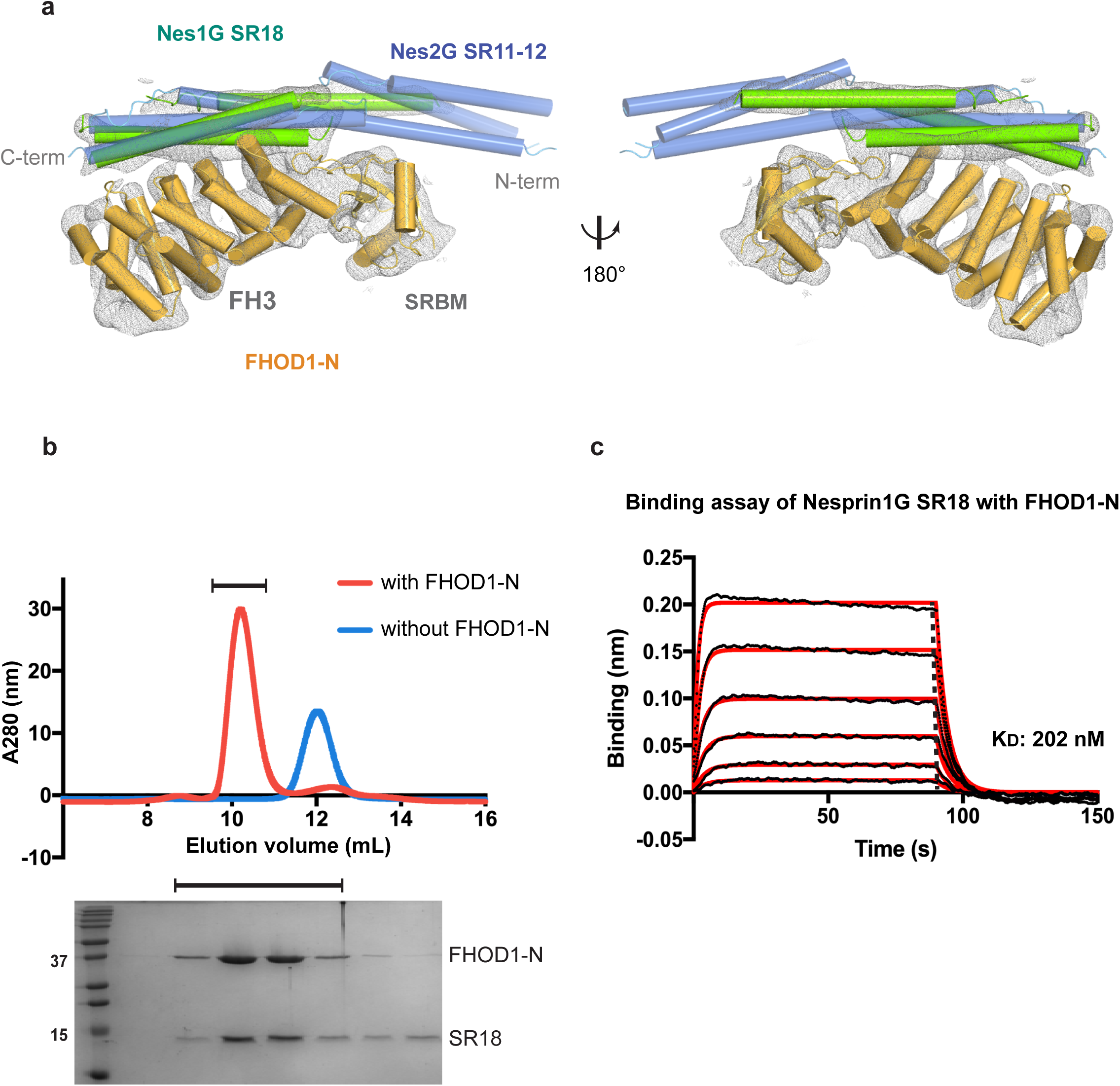
SR18 of Nes1G is sufficient to bind FHOD1-N. **a**, Crystal structure of FHOD1-N bound with Nes1G-SR17-18 at 7.0 Å resolution with 2Fo-Fc map contoured at 1.2 sigma. SR17 was not modeled because of insufficient electron density. Nes2G-SR11/12 (blue) overlaid for comparison. **b**, SEC experiment of Nes1G-SR18 with or without FHOD1-N, accompanied by SDS-PAGE analysis of the eluted fractions. **c**, Binding of Nes1G SR18 with FHOD1-N measured by BLI.

### Binding of Nes2G does not preclude FHOD1-autoinhibition

Actin bundling by FHOD1-N is autoinhibited by its C-terminal DAD-helix binding to the FH3 domain. This autoinhibition can be released by phosphorylation of various serine and threonine residues within the DAD-helix by Rho dependent kinase^24^. Since both, Nes2G-SR11/12 and DAD, can bind to the FH3 domain, we tested whether they compete with one another. While the DAD binding site is not structurally known for FHOD1, it can be predicted with great confidence. The superposition of the FH3 domain of mDia bound to its DAD-helix (PDB code 2F31) onto FHOD1-FH3 suggests a position, that is functionally confirmed by mutational analysis. A V228E mutation on the surface of FHOD1-FH3, predicted to be in the core of the DAD-binding site, potently blocks DAD-binding *in vitro*^20^. Modelling a putatively bound DAD-helix onto our Nes2G- SR11/12-FHOD1-N complex suggests that the two interactions are independent and not in steric conflict (Figure 7a). To test this experimentally we co-incubated Nes2G-SR11/12-FHOD1- N complex with an MBP-tagged 3C-cleavable DAD-helix fusion. To avoid steric hindrance by the MBP-fusion tag we added 3C protease. As expected, we detect DAD-helix coeluting with the Nes2G-SR11/12-FHOD1-N complex, indicating that the two binding sites are independent (Figure 7b). It further suggests that FHOD1 autoinhibition does not regulate Nes1G or Nes2G binding^25,26^

**Figure 7:**
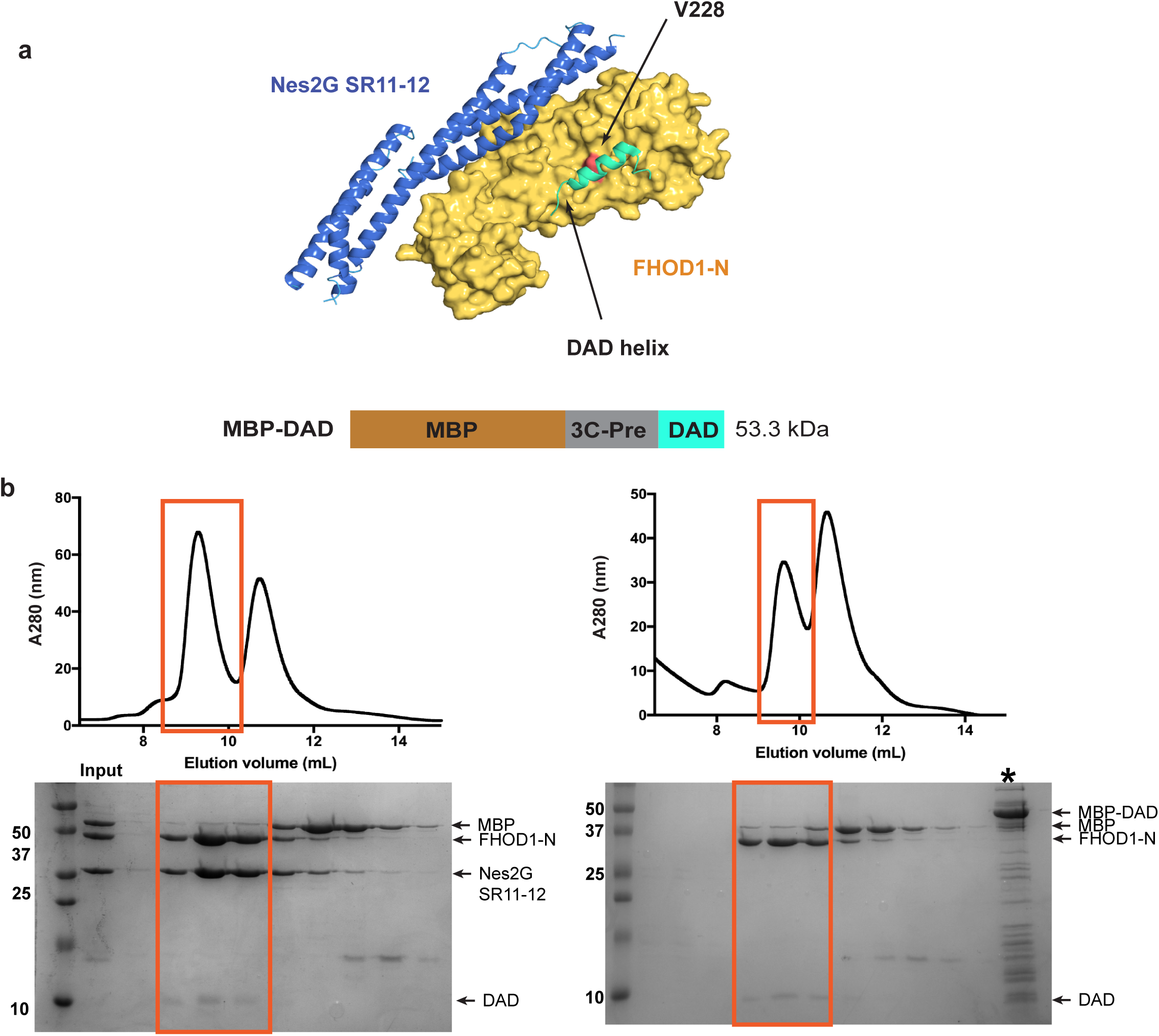
SR binding to FHOD1 does not sterically compete with DAD autoregulatory binding. **a**, The Nes2G-SR11/12-FHOD1-N complex overlaid with the DAD helix (cyan). The DAD helix was positioned based on an alignment with the mDia-FH3 domain bound to its DAD helix (PDB code 2F31). FHOD1-V228, critical for DAD-binding in deep-orange. **b**, Analytical SEC of MBP- DAD with Nes2G-SR11/12 and FHOD1-N complex (left panel) or FHOD1-N alone (right panel) in the presence of 3C protease. A Superdex S75 10/300 GL column was used in all the SEC experiments. Uncleaved MBP-DAD (*) was run as a control.

### FHOD1 has a spectrin-repeat binding modulator (SRBM) rather than a GTPase-binding domain

Phylogenetic analysis revealed that the FH3 or DID domain in formins is preceded by one of two small domains. either a GTPase-binding domain (GBD or G) or the SRBM described here (formerly GBD2 or G2). Several crystal structures of the tethered GBD-FH3 domain in complex with small GTPases exist from diverse species, and the binding mode is very similar. Both the GBD domain and the FH3 domain contribute to the GTPase interaction (Figure 8a)^18,19^. In comparison, the tethered SRBM-FH3 domain instead binds SRs, also with contributions from both domain elements (Figure 8b). When we phylogenetically analyzed maximally diverged SRBM-FH3 sequences and compared them with equally diverged GBD-FH3 sequences we detected distinct conservation of key residues within the respective binding sites of the FH3 domain (Figure 8). For example, positions 144 and 189 of FHOD1, strongly conserved as glutamine and leucine, respectively, in the SRBM-FH3 family, are instead strongly conserved as valine and asparagine in the GBD-FH3 family (Supplementary fig. 3D). Conversely, the pair of invariant arginines in position 136 and 137 of FHOD1, important for SR recognition, are not at all conserved in the GBD-FH3 family (Supplementary fig. 3H). This result clearly suggests that the FH3 domain has evolved to either bind a small GTPase or SRs, depending on which domain, SRBM or G, it is tethered to. In other words, SRBM or G modulate the binding activity of the neighboring FH3 domain.

**Figure 8:**
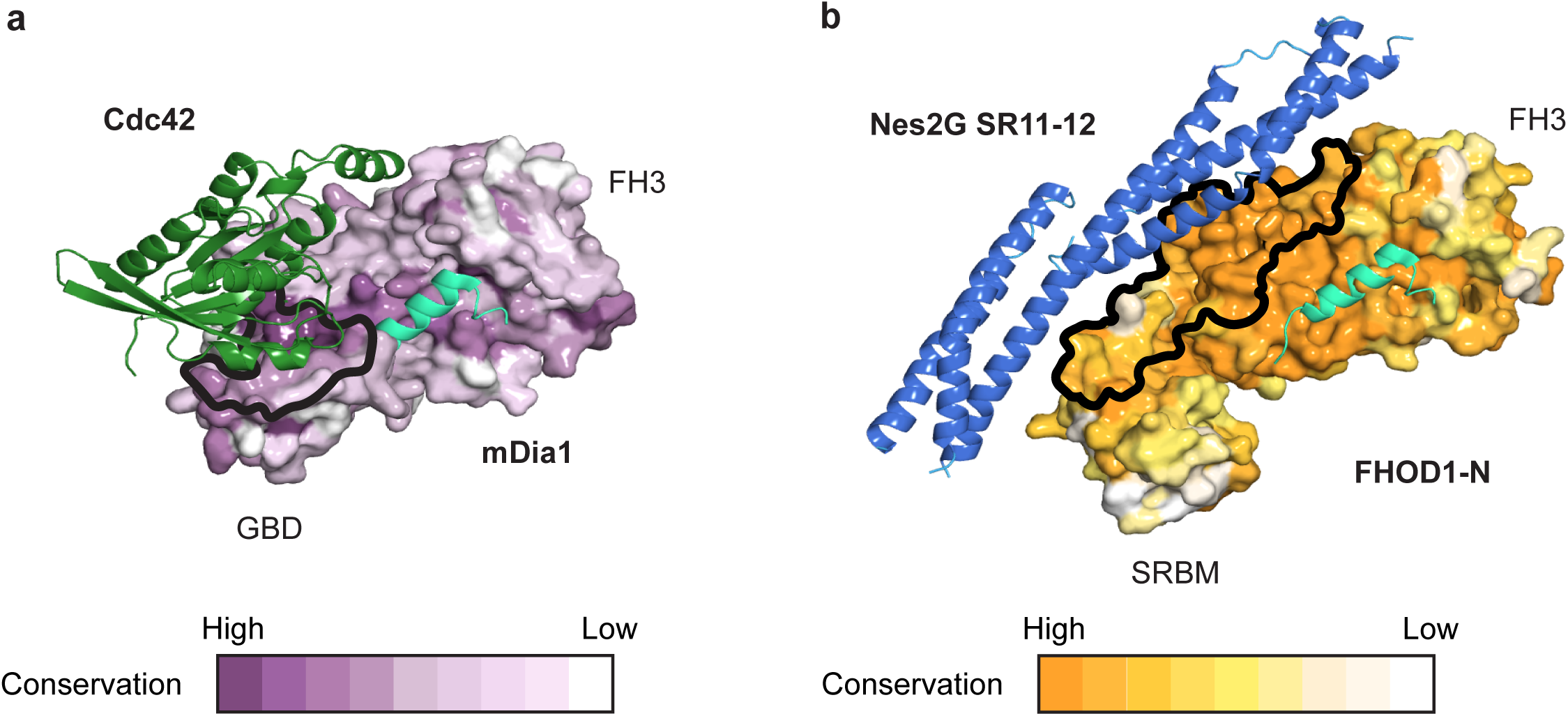
Modulation of the FH3 binding activity by its N-terminally paired domain. **a**, Comparison between GBD-FH3 of mDia1 bound to the small GTPase Cdc42 and **b**, SRBM- FH3 of FHOD1 bound to Nes2G-SR11/12. Structures are aligned using the FH3 domain. Binding surfaces outlined. Paired SRBM- or GBD-FH3 domains shown as surfaces, gradient-colored according to conservation as indicated. DAD helix position modeled (cyan). SR and small GTPase bind in mutually exclusive positions. Surfaces are specifically conserved to recognize either SR or small GTPases, defining a difference between the two classes of formins.

## DISCUSSION

Here we show how FHOD1-N engages with SRs from two nesprins to form stable complexes. How does this structure inform us about tethering between the nucleus and actin bundles during nuclear migration along TAN lines? One important factor is how to achieve stability within this protein network. We propose that this is established through a multitude of interactions with actin. Specifically, Nesprin2 has two N-terminal CH domains each of which directly bind actin filaments. Furthermore, SR11/12 engages with FHOD1, which bundles actin through its FH2 domain. Since FHOD1 itself is a dimer, this tethered FHOD1-Nes2G heterotetrameric unit then already has 6 individual actin binding sites (Figure 9a). Due to avidity, this arrangement should greatly increase actin filament affinity, a prerequisite for sustaining the large mechanical forces associated with moving such a gigantic cargo. This is also compatible with a model of Nesprins1/2 tethering together neighboring actin filaments to induce bundling in a TAN line assembly (Figure 9b)^27^.

**Figure 9:**
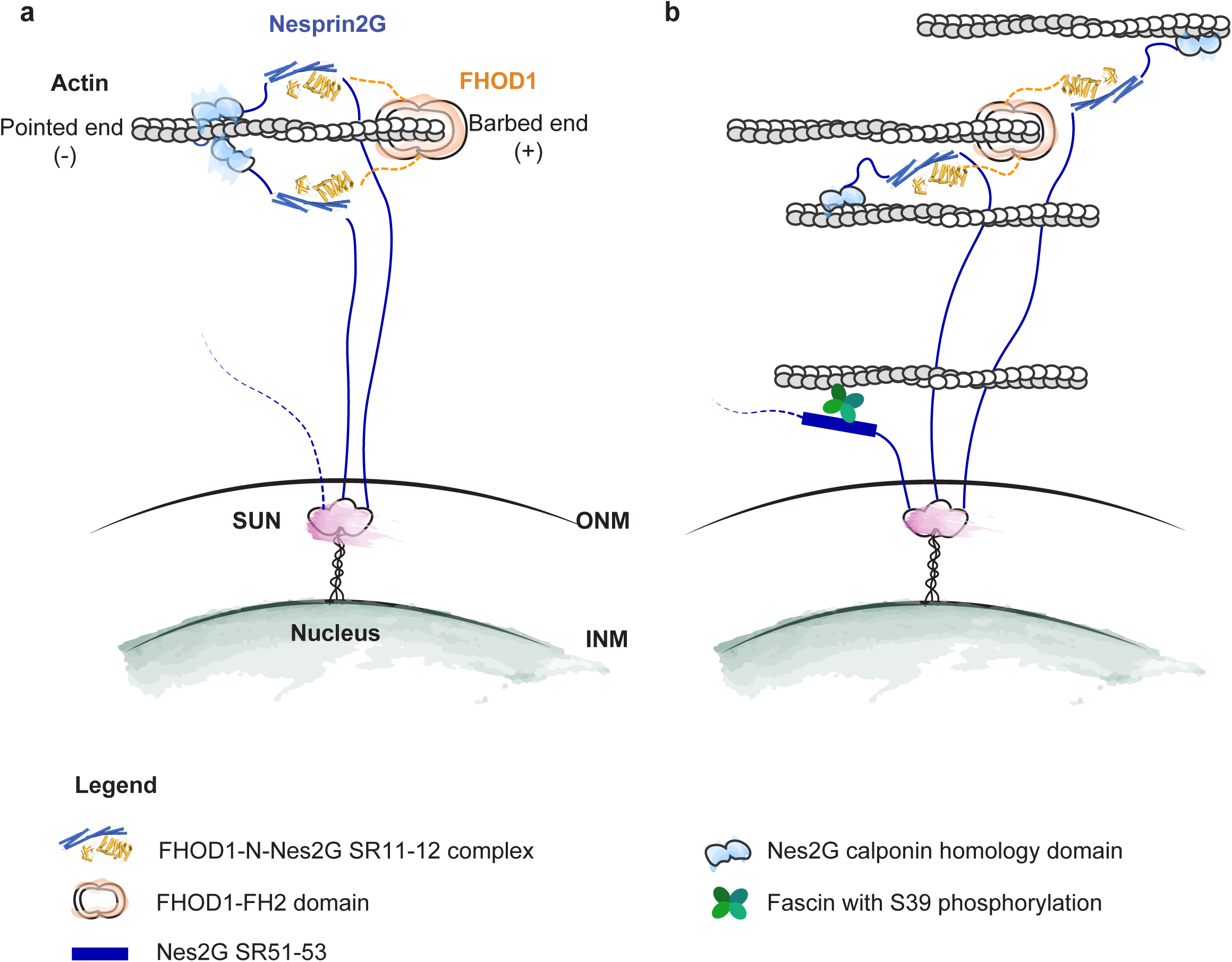
Model for connecting actin bundles with the nucleus. A cartoon depiction of the Nes2G-FHOD1 structure in the context of TAN lines. FHOD1 binds to the barbed end of long actin polymer. **a**, Its association with Nes2G can either enforce binding of LINC to a single actin polymer or **b**, could play a part in the bundling of actin cables.

This structure is a snapshot along the path toward understanding the entire TAN line network. Much remains to be discovered to mechanistically understand the whole process of nuclear migration. One important element to consider going forward is the significance of FHOD1 binding to primarily just one SR in both, Nes1G and Nes2G. With these giant nesprins containing 74 and 56 SRs, respectively, is there significance in the position of the interacting SR? Going forward, an interesting experiment will be to shuffle the position of the FHOD1 binding site within these long SR proteins and see which effect this may have. The existing data strongly suggest that the position is critical. Our structures reveal that FHOD1 binds only highly specific SRs, namely those with the conserved sequence motif we discovered. It explicitly does not bind generic SRs. Therefore, the position of the cognate SR within a nesprin should matter. An alternative hypothesis would be that the number of binding sites rather than the position within a nesprin is important. This is also a testable scenario.

This is the first crystal structure of a nesprin SR domain and it confirms that Nes2G-SR11/12 shares, as expected from modeling, strong structural homology with the canonical SR architecture, as seen in α- and β-spectrin (Supplementary Fig.5A-B). The main interaction with FHOD1 is mediated through a binding interface which has a distinct sequence motif DxWLD[IVLA]xE, found in Nes2G-SR12 and Nes1G-SR18, respectively. Binding by Nes1G and Nes2G is very similar, except that for Nes2G the interface also includes a short element of the preceding SR11. For Nes1G, SR17 is dispensable. While one could formally argue that in Nes2G a tandem SR is recognized while in Nes1G a single SR is sufficient, we would consider this to be an exaggeration of the actual differences. The SR11 element bound by FHOD1 is merely through the shared central helix that connects SR11 and SR12 and defining the exact domain boundary between the two is to an extend somewhat arbitrary. To our knowledge, there is one crystal structure of SRs with a binding partner, which is human erythroid β-spectrin bound to ankyrin (PDB code 3KBT)^28^. Here, the interface clearly spans two SRs and the authors argued for the specific importance of the relative orientation of the neighboring domains. The binding itself is very different from the FHOD1-SR interface, which is not surprising given that FHOD1 and ankyrin have different domain architectures.

We now establish that the formerly named G2 or GBD2 domain of FHOD1 is a spectrin-repeat binding modulator, which we call SRBM. It is another remarkable example of how nature has coopted a common domain fold, here ubiquitin, to evolve a novel function. This common theme has been seen in many other folds, including helix-turn-helix motifs, β-propeller proteins, to name just a few ^29,30^. The distinct evolution of the SRBM within the larger formin family has been recognized before based on phylogenetic analysis^31^. However, it remained nearly unchallenged that this domain may not bind a small GTPase. Our work provides another powerful example for why experimental validation is a critical element of discovery. Divergent evolution of domains into diverse functions can be difficult to discern, especially if these functions do not correlate with an easily recognizable sequence pattern.

The SRBM is not an autonomous domain that has its separable function. It appears to only function in tandem with the neighboring FH3 domain, which is why we call it a modulator. Neither FH3, nor SRBM bind a spectrin repeat independently. The sequence conservation of the SRBM supports this notion. The best conserved part within the SRBM is the long loop connecting β1 and β2, which is precisely necessary to lock the module with the FH3 domain and also to directly bind the cognate SR. Furthermore, neighboring SRBM and FH3 domains coevolved to bind spectrin repeats, which is evident when one compares the FHOD class of formins (Figure 8). The situation is not very different for the bona fide GBD, which is found in the larger class of diaphanous-related formins (DRFs). As shown previously, it also binds small GTPases in conjunction with the neighboring FH3 and apparently not autonomously^18,19^. In the DRFs, the GBD co-evolved with the neighboring FH3 in much the same way as SRBM co-evolved with its joint FH3. This is clearly evident in the Pfam protein family database, where the GBD is defined as containing about half of the FH3 domain (Pfam entry 06371), with conserved residues distributed over both structural elements. Taken together, we suggest that GBD and SRBM are two, mutually exclusive modulators of the FH3 domain that direct and specify its binding activity. Perhaps, it is no coincidence that the FH3 domain is an α-solenoid, a versatile domain found in many, highly diverse protein assemblies.

## MATERIALS & METHODS

### Protein expression and purification

Residues 1425-1649 of human Nesprin-2, corresponding to spectrin repeats 11 and 12, were cloned into the bacterial expression vector pETDuet1 to generate a human rhinovirus 3C (3C)- cleavable His-tag fusion protein (Nes2G-SR11/12). LOBSTR(DE3)-RIL *E. coli* cells (Kerafast) were transformed with the expression vector and grown at 37 °C in LB medium with 0.4% (w/v) glucose and in the presence of antibiotics to maintain the plasmids. Protein expression was induced with 0.2 mM isopropyl β D-1-thiogalactopyranoside (IPTG) at OD600 = 0.7, at which point the culture was shifted to 18 °C. Cells were grown for another 12-16 h. The cells were then harvested by centrifugation at 6000 g for 6 mins and resuspend in 50 mM potassium phosphate pH 8.0, 400 mM NaCl, 40 mM imidazole and 5 mM β-mercaptoethanol. Cells were lysed by mechanical disruption with a LM-20 Microfluidizer (Microfluidics) at 18,000 psi then spun at 9500 g for 25 mins. The clear supernatant was incubated with Ni-Sepharose 6 Fast Flow (GE Healthcare) to capture the His-tagged protein. After batch washing in lysis buffer, the Ni-resin was poured into a disposable column, drained, and the bound protein was eluted with elution buffer (10 mM Tris/HCl pH 7.5, 150 mM NaCl, 250 mM imidazole, 5 mM β-mercaptoethanol). The eluate was cleaved with 3C protease overnight and Nes2G-SR11/12 was further purified by cation exchange chromatography on a HiTrap SP-FF column (GE Healthcare).

Similarly, the coding sequence for residues 1-339 of human FHOD1 was also cloned into the pETDuet1 plasmid (wtFHOD1-N), as well as a C31S C71S double mutant (mtFHOD1-N). Expression and purification were carried out as for Nes2G-SR11/12 described above. To form complexes, purified Nes2G-SR11/12 and wt FHOD1-N or mt FHOD1-N were mixed in a 1 to 1 ratio followed by size exclusion chromatography. The protein complex eluted as a monodisperse peak from a Superdex 200 gel filtration column (GE Healthcare) in 20 mM HEPES/NaOH pH 8.0, 100 mM KCl, 0.2 mM EDTA, 1 mM DTT. All remaining Nes2G-SR11/12 mutants (L1628R, L1628A E1632A, D1625A D1629A) and FHOD1-N mutants (R136A R137A, E36A P37A R38A) were expressed and purified with the same protocol.

FHOD1 DAD domain (residue 1052-1164) preceded by maltose binding protein (MBP) and a 3C-cleavage site was expressed the same way. The sample in lysis buffer 10 mM HEPES/NaOH pH 8.0, 200 mM KCl, 5mM β-mercaptoethanol was purified by affinity chromatography over amylose resin. The target protein was eluted with 10 mM maltose and further purified via gel filtration on a Superdex S75 (GE Healthcare) column.

Three Nesprin1 constructs were generated; The first containing residue 1976 to 2200 and identified as SR17/18 (Nes1G-SR17/18); the second a D2176A D2180A mutant and the third an SR18 only construct (residue 2076-2200). All were cloned, expressed, and purified following the method described above without substantive changes. For affinity measurements by bio- layer interferometry, Nes1G an Nes2G constructs were N-terminally tagged with an AVI sequence (GLNDIFEAQKIEWHE) for biotinylation in the *E. coli* expression host.

### Crystallization

Purified complexes of Nes2G-SR11/12 with wtFHOD1-N or mutant FHOD1-N were concentrated to 7 mg/ml for sparse matrix crystallization screen. Both complexes initially crystallized in 0.1 M Tris/HCl pH 8.5, 0.2 M potassium sodium tartrate, 21% PEG 3350 at 18 °C using the hanging-drop vapor diffusion method with a 1 μL well-solution to 1 μL protein setup. Large, rectangular-rod crystals formed within 5 days and were supplemented with 15% glycerol as cryo-protectant prior to freezing. Crystals of Nes1G-SR17/18 and FHOD1-N were grown in 0.1 M HEPES/NaOH pH 7.0, 0.2 M sodium thiocyanate, 40% 5/4 pentaerythritol propoxylate at 18 °C using the sitting drop method.

### Data collection & processing

Diffraction data were collected at the NE-CAT beamline 24-ID-C in Argonne National Laboratory and processed with HKL-2000^32^. The structures were all solved by molecular replacement using Phaser from within the PHENIX package^33^. The initial search model was FHOD1-N (PDB Code 3DAD)^34^. Additional helical density for the spectrin repeats was readily visible in the initial map. The model was built and refined iteratively using COOT^35^ in combination with phenix.refine. The structure of Nes1G-SR17/18 in complex with wtFHOD1-N were phased with molecular replacement of the Nes2G_-_SR11/12 FHOD1-N complex structure. Table 1 lists the final statistics for two complexes, Nes2G-SR11/12 - FHOD1-N and Nes1G-SR17/18 - FHOD-N respectively.

### Complex analysis by size exclusion chromatography

Purified proteins were mixed in a 2 to 1 molar ratio, with Nes1G or Nes2G in excess, and incubated on ice for 1hr. Then, 1 mL of sample mixture was loaded onto a Superdex S200 10/300 GL gel filtration column for size analysis. The running buffer for all analytical studies was 20 mM HEPES/NaOH pH 8.0, 100 mM KCl, 0.2 mM EDTA, 1 mM DTT. The chromatography experiment was run at 0.5 ml/min flow rate and 500 μL sample fractions were collected for SDS-PAGE analysis.

### Bio-layer interferometry

Interaction of Nes1G or Nes2G spectrin repeats with FHOD1 was measured by Bio-layer interferometry on an 8-channel Octet RED96e system (Forté Bio). First, streptavidin biosensor tips were pre-incubated with an assay buffer (20 mM HEPES/NaOH pH 8.0, 100 mM KCl, 0.2 mM EDTA, 1 mM DTT, 0.2% bovine serum albumin (BSA) and 0.01% Tween-20) for 10 min at 30 °C. Then, the tips were incubated with N-terminally biotinylated Nes1G and Nes2G constructs in assay buffer to yield a loading thickness of 0.3-0.4 nm. After washing the tips with assay buffer binding to wtFHOD1-N was measured in real time by recording the increase in optical thickness of the tips during an association phase. Finally, the tips were transfer back into assay buffer to measure the dissociation rate. The concentration of FHOD1 was fixed at 0.4 μM for qualitative experiment to validate the effect of mutant versus wild-type Nes1G and Nes2G binding. A two-fold dilution series of FHOD1-N concentration ranging from 0.01 to 0.4 μM was used for measuring the binding affinity of FHOD1 and Nesprins. For data analysis we used Octet Data Analysis software and transferred the processed output into GraphPad Prism 7 for curve-fitting analysis by using association-dissociation non-linear regression model.

### DAD helix binding with Nes2G and FHOD1-N

Purified MBP-DAD was mixed with either wtFHOD1-N only or Nes2G-SR11/12-FHOD1-N complex in a 1 to 1 molar ratio and incubated overnight in the presence of 3C protease. The samples were analyzed by gel filtration over Superdex S75 10/300 in running buffer 20 mM HEPES/NaOH pH 8.0, 100 mM KCl, 0.2 mM EDTA and 1 mM DTT. Relevant elution fractions were analyzed by SDS PAGE.

### Analytical ultracentrifugation

FHOD1-N (2 mg/mL) was dialyzed into 20 mM HEPES/NaOH pH 8.0, 100 mM KCl, 0.2 mM EDTA with and without 0.5 mM Tris (2-carboxyethyl) phosphine hydrochloride (TCEP-HCl) for two days in 4°C. Samples were then prepared in 450μL with final concentrations of 0.3 mg/mL, 0.6 mg/mL and 0.8 mg/mL for sedimentation velocity (SV) experiment. The experiment was performed in an Optimal XL1 (Beckman Coulter) ultracentrifuge using the An50 Ti rotor at 20 °C with a rotation speed of 45,000 rpm. A total of 500 absorbance scans at 280nm were acquired with 1-minute intervals. Scans 1 to 50 were used for analysis in SEDFIT ^36^.using the size distribution c(s) model. SEDNTERP were used to calculate the density and viscosity of the buffer at 20°C.

### Plasmids for LPA stimulation experiments

All constructs were confirmed by DNA sequencing. pSUPER-hygro was derived from the pSUPER-puro (Oligoengine) by replacing a puromycin resistance gene sequence with a hygromycin resistance gene sequence from pMSCV-hygro vector (Clontech), and it was used for expressing shRNA in NIH3T3 fibroblasts by retroviral infection. pMSCV-puro EGFP-C4 vector was used to express EGFP tagged protein in NIH3T3 by retroviral infection. Both pMSCV-puro EGFP-C4 human FHOD1 WT and I705A mutants were previously described ^23^. Human FHOD1 R136A R137A mutant was made by introducing (CGCCGC to GCCGCC) point mutation. The FHOD1 cDNA was inserted into pMSCV-puro EGFP-C4 vector with BamHI and NotI restriction sites. The shRNA sequences for NC and FHOD1 were 5’- caacaagatgaagagcaccaa-3’ and 5’-gaacctctttcctaccatttc-3’, respectively.

### Cell culture, virus production, infection, and drug selection

NIH3T3 fibroblasts were maintained in Dulbecco’s Modified Eagle Medium (DMEM; Corning Inc.) containing 10 mM HEPES pH 7.4 and 10% (v/v) bovine calf serum (GE Health Life Science). Meanwhile, 293T cells were maintained in DMEM containing 10 mM HEPES pH 7.4 and 5% (v/v) bovine calf serum and 5% (v/v) fetal bovine serum (Gemini Bio-Products). The 293T cells were transfected with retroviral vectors and pantropic packaging plasmids. Medium containing the produced virus was harvested 24 hr after transfection, added to the NIH3T3 fibroblasts in the presence of 2 μg/ml polybrene (EDM Millipore TR_1003) and incubated for one day. The infected cells were selected with 300 μg/ml hygromycin B (EDM Millipore 400051) and 1.5 μg/ml puromycin (Sigma-Aldrich P8833) for one week. The cells were recovered from drug treatment for one additional week before used for the experiments.

### Immunofluorescence microscopy

For indirect immunofluorescence microscopy, cells were fixed with 4% paraformaldehyde in phosphate-buffered saline (PBS) for 20 min, then they were permeabilized and blocked with PBS containing 0.1% Triton-X and 1% BSA for 30 min. The cells were labeled first with primary antibodies and then fluorescently-labeled secondary antibodies (EMD Millipore AB16901) and DAPI (Thermo Fisher Scientific D3571). Images were acquired with a 40× PlanApo objective (NA 1.0) and DS-Qi2 (Nikon) on a Nikon Eclipse Ti microscope controlled by Nikon’s NIS-Elements software.

### Western blotting

Proteins were separated by SDS-PAGE then transferred onto a nitrocellulose blots, probed with antibodies and detected either by chemiluminescence with Odyssey Fc (LI-COR Inc.) or infra- red fluorescence with Odyssey CLx (LI-COR Inc.). The antibodies used were mouse monoclonal GFP antibody (Santa Cruz, sc-9996), rabbit polyclonal FHOD1 antibody (Santa Cruz, sc-99209, discontinued, rat Monoclonal tyrosinated-alpha-tubulin (YL-1/2) (European Collection of Authenticated Cell Cultures, 92092402).

### LPA stimulation

A day before serum-starvation, NIH3T3 fibroblasts were plated on acid-washed coverslips. The next day, cells at about 40% confluency on coverslips were washed three times with DMEM and then DMEM containing 10 mM HEPES pH 7.4 and 0.1% (v/v) fatty acid free bovine serum albumin (Sigma-Aldrich, A7906) was added. After two days serum-starvation, the cells were stimulated with 10 μM LPA (Avanti Polar Lipids, 857130P) and fixed after 2 hours.

### Cell analysis

The position of centrosome relative to the axis between the nuclei and the leading edge was analyzed from images of DAPI, tubulin and β-catenin/pericentrin antibody-labeled cells as previously described ^37,38^. Nuclear and centrosomal positions of NIH3T3 fibroblasts were determined from images using Cell Plot software ^39^. Statistical analysis of data on centrosome reorientation was assessed by Chi-square test using GraphPad Software. Statistical evaluation of the position of the nucleus and centrosome in NIH3T3 fibroblasts was by one-way ANOVA followed by Tukey’s multiple comparison test using SAS. All evaluated data were from at least N=3 experiments.

## Acknowledgements

Research was supported by the US NIH under grant number R01-AR065484 (T.U.S.) and T32GM007287 (V.C.). The X-ray crystallography work was conducted at the APS NE-CAT beamlines, which are supported by NIH award GM103403. Use of the APS is supported by the US Department of Energy, Office of Basic Energy Sciences, under contract no. DE-AC02- 06CH11357.

## Author Contributions

S.M.L. and T.U.S. designed the study. S.M.L. and V.E.C. performed the experiments. G.G.G. and S.A. contributed Figure 4. S.M. L. and T.U.S. interpreted the results and wrote the manuscript with input from G.G.G., S.A., and V.E.C..

