## Supplementary Material for "Structures of FHOD1-Nesprin1/2 complexes reveal alternate binding modes for the FH3 domain of formins"

#### Supplementary Figure 1

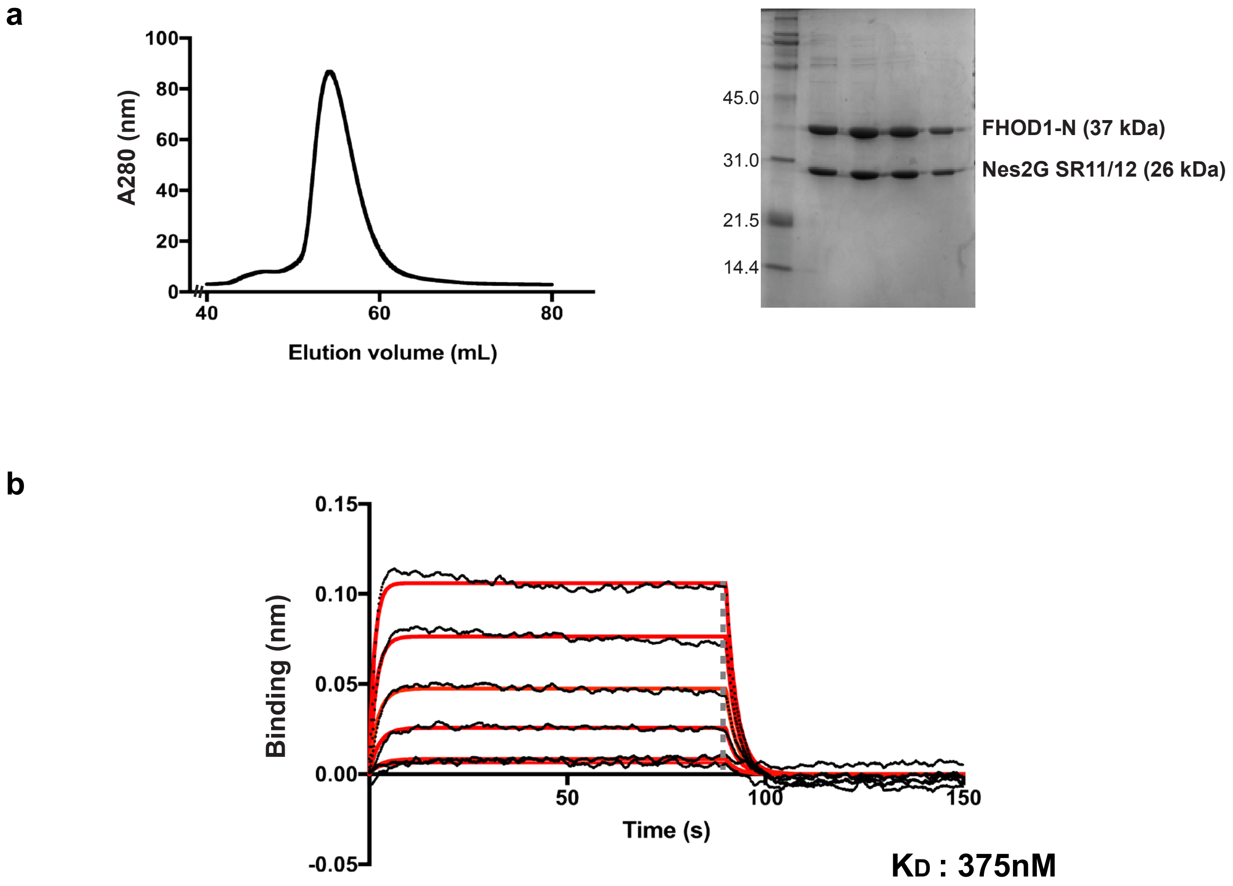


**Supplementrary Figure 1:** Purification and binding analysis of Nes2G SR11-12 and FHOD1 complex.
**a,** Size exclusion profile and corresponding SDS PAGE gel of FHOD1-N in complex with Nes2G in 20 mM HEPES pH 8.0, 100 mM KCl, 0.2 mM EDTA,
1 mM DTT using a Superdex75 16/60 column. **b,** BLI binding profile of biotinylated Nes2G SR11-12 with FHOD1-N, concentration of FHOD1-N ranges from 25 - 400 nM in 2-fold serial dilution. Binding buffer is 20 mM HEPES pH 8.0, 100 mM KCl, 0.2 mM EDTA, 1 mM DTT, 0.2 % BSA and 0.01 % Tween 20. Red lines represent fitted curves.

### Supplementary Figure 2

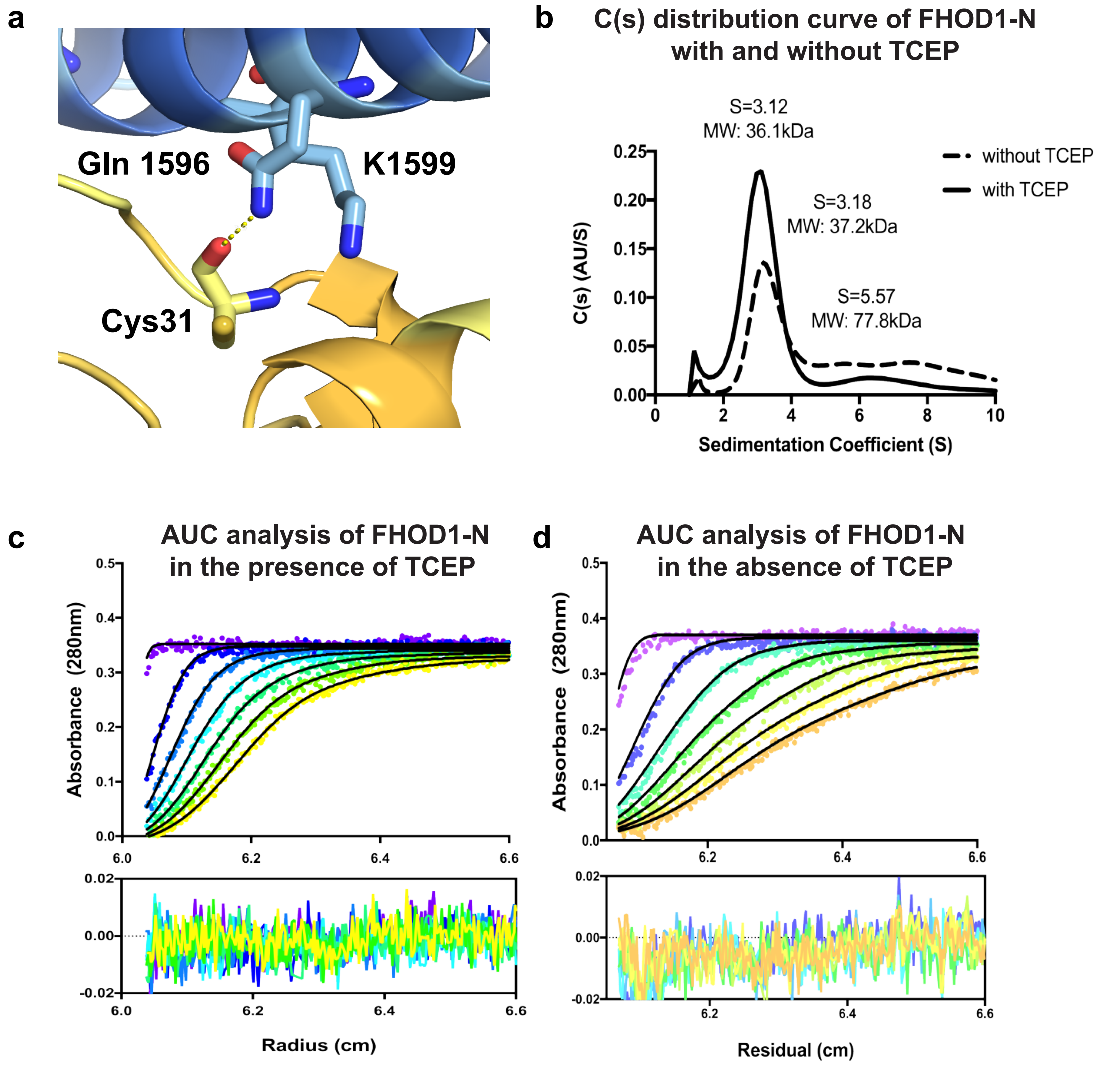


**Supplementary Figure 2**: Cys 31 and Cys 71 of FHOD1-N are not specifically involved in complex binding with Nes2G-SR11-12.
**a.** Non-specific interaction of Cys 31 backbone with Nes2G-SR11-12 **b.** The c(s) distribution profile of wtFHOD1-N in the presence (solid line) and absence of TCEP (dotted line) with corresponding sedimentation coefficients and molecular weights for the species present in the samples. **c,** Representative radial absorbance profiles at 280 nm as a function of time for 0.25 mM FHOD1-N dialyzed in 20 mM HEPES pH 8.0, 100 mM KCl, 0.2 mM EDTA with or **d,** without 0.5 mM TCEP. Residual for the fits are shown in the lower panels.

#### Supplementary Figure 3

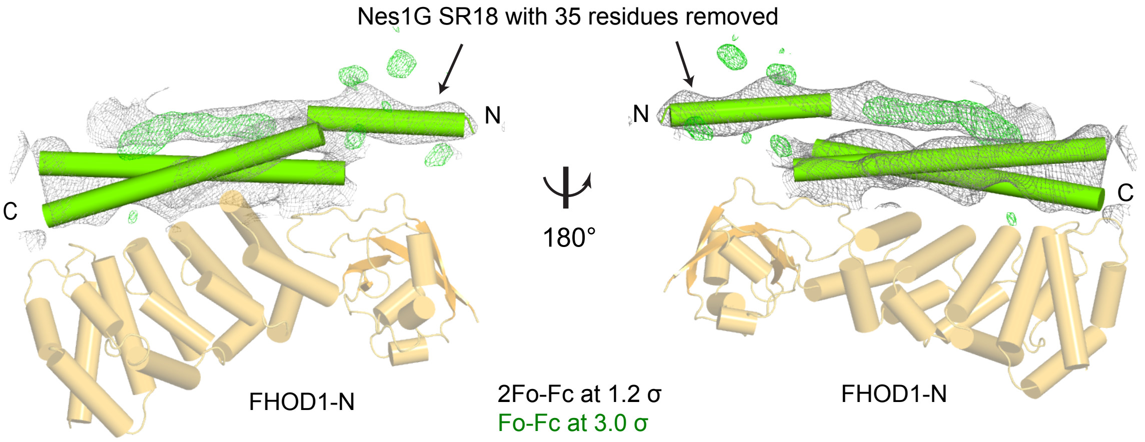


**Supplementary Figure 3:** Validation of molecular replacement solution for Nes1G-SR17/18 - FHOD1-N complex. We calculated 2Fo-Fc (grey) and difference density Fo-Fc (green, 3σ) after removing a 35-residues helical segment from Nes1G-SR18 model. The difference density indicates that the solution is correct.

#### Supplementary Figure 4

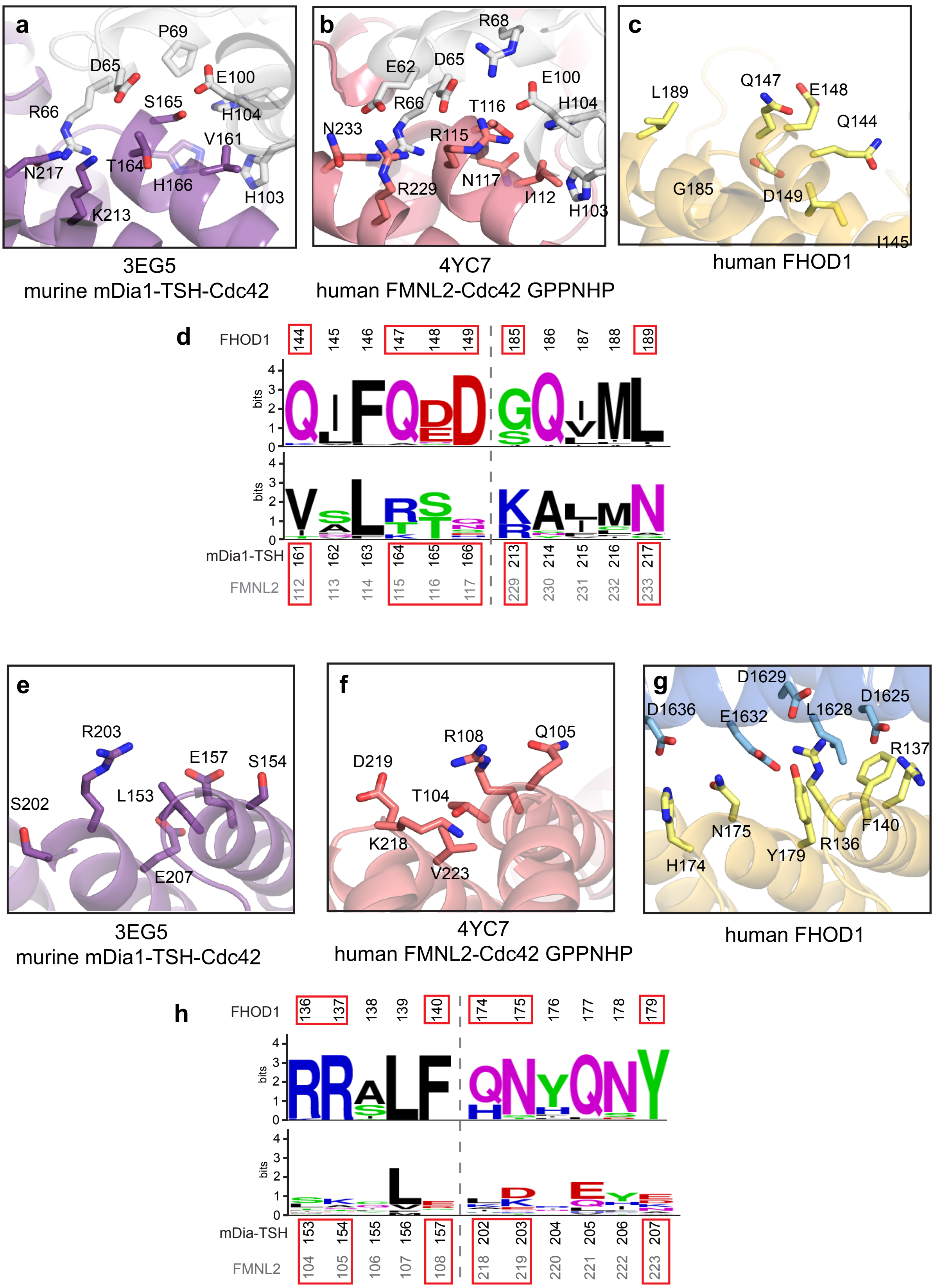


**Supplementary Figure 4**: Comparison of the FH3 domains between Drf family members and FHOD1 in the context of GTPase vs. SR binding.
**a,** Residues of the GTPase binding site for both mDia and **b,** FMNL2 are shown as sticks. The GTPase is colored in grey. **c,** The corresponding residues on FHOD1 based on FH3 structural alignment. **d,** Weblogo showing the conservation of these residues among GTPase binding Drf members in comparison with FHOD1/3 members; residues shown in panels **a-c** are boxed in red. **e-g,** Vice-versa, residues at the SR binding site for FHOD1 are represented as sticks in **g** with the structurally equivalent positions for mDia in **f** and FMNL2 in **g**. **h,** Weblogo focus on the SR-binding positions important for FHOD1 (boxed red) and the equivalent positions in Drfs.

#### Supplementary Figure 5

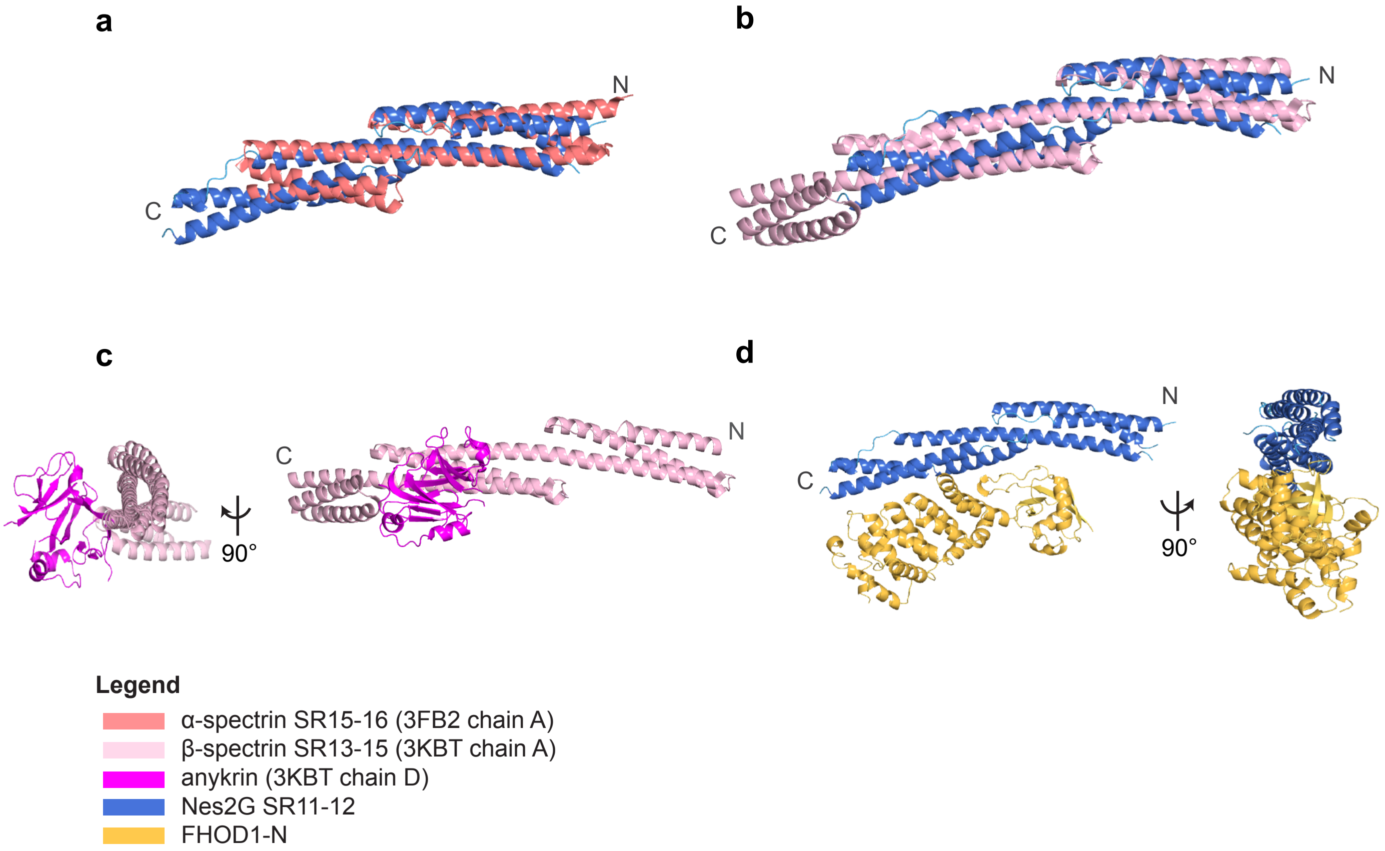


**Supplementary Figure 5**: Structural compariosn of spectrin repeats and their interactors. **a,** Superposition of Nes2G-SR11/12 with human brain α-spectrin and **b,** erythroid β-spectrin show that Nes2G SR11-12 has a canonical SR architecture with connected three-helix bundles. **c,** Cartoon representation of β-spectrin in complex with ankyrin in the same perspective as **d,** Nes2G11/12-FHOD1 complex.
